## Supplementary file for "Impact of dendritic non-linearities on the computational capabilities of neurons"

### CONTENTS

A. Analytical results 1
1. Definition of the non-linear model of the neuron 1
2. Training set and partition function 2
3. Replica method 2
4. Replica Symmetric analysis 4
a. Large $K$ limit 4
b. Free entropy and saddle point equations 7
c. Effective order parameters for some non-linearities 7
d. Small data regime 9
e. Distribution of dendritic preactivations 9
f. Limit of large somatic thresholds 9
5. Critical capacity 10
a. Limit of large dendritic threshold 12
b. Limit of small dendritic threshold 12
6. Distribution of synaptic weights 12
a. Distribution of synaptic weights in the maximal storage limit 13

B. Numerical experiments 15
1. Choice of the thresholds 15
2. Choice and scaling of the hyper-parameters 17

References 18

### Appendix A: Analytical results

#### 1. Definition of the non-linear model of the neuron

We recall here the main definitions of the single neuron model studied in the main text of the paper. Given an activity pattern $\boldsymbol{\xi}^\mu \in \{0,1\}^N$, the output of our model of neuron is obtained in two steps. Firstly, the activity pattern is processed by the corresponding dendritic branch; we suppose here that we have $l = 1, \dots, N/K$ dendritic branches each having a set of $i = 1, \dots, N$ positive synaptic weights $W_{li}$. The output activity $\tau_l^\mu$ of a given branch $l$ that corresponds to the activity pattern $\boldsymbol{\xi}^\mu$ is obtained as

$$\tau_l^\mu = g\left(\sqrt{\frac{K}{N}}\sum_{i=1}^{N/K} W_{li}\xi_{li}^\mu - \sqrt{\frac{N}{K}}\theta_d\right) \equiv g\left(\lambda^\mu\right) \tag{A1}$$

where $\theta_d$ is a threshold modeling inhibition at the level of the dendritic branch, while $g(\cdot)$ is a generic positive, (possibly) non-linear function. Secondly, the output of each branch is combined linearly by using another set of $K$ synaptic weights $c_l$, $l = 1, \dots, K$ and the output is obtained as

$$\sigma^\mu_{\text{out}} = \Theta\left[\frac{1}{\sqrt{K}}\sum_{l=1}^{K} c_l\,\tau^\mu_l - \sqrt{K}\theta_s\right] \tag{A2}$$

where $\Theta(x)$ is the Heaviside theta function that is 1 if $x > 0$ and 0 otherwise. The parameter $\theta_s$ is a threshold modelling inhibition coming from inhibitory neurons. In the following we will consider, for simplicity $c_l = 1$ for every $l = 1, \dots, K$.

#### 2. Training set and partition function

We consider a training set composed of $P = \alpha N$ random i.i.d. activity patterns $\boldsymbol{\xi}^\mu \in \{0,1\}^N$ and i.i.d. labels $\sigma^\mu \in \{0,1\}$ with $\mu = 1, \dots, P$. The probability distribution of each component of a pattern is given by

$$P(\xi^\mu_{li}) = f_{\text{in}}\,\delta(\xi^\mu_{li} - 1) + (1 - f_{\text{in}})\,\delta(\xi^\mu_{li}) \tag{A3}$$

where $f_{\text{in}}$ is the *input coding level* of the patterns. We consider a probability distribution of labels to be equal in form to (A3) but we allow the possibility to have different coding level in the output $f_{\text{out}}$.

In order to study the volume of synaptic weights that correctly associate to a given pattern of activity $\boldsymbol{\xi}^\mu$ the corresponding label $\sigma^\mu$ we use a standard statistical mechanics approach [1, 2]. Firstly, we define the characteristic function

$$X_{\xi,\sigma}(W) = \prod_\mu \Theta\left(\frac{(2\sigma^\mu - 1)}{\sqrt{K}}\left(\sum_{l=1}^{K} c_l\,\tau^\mu_l - K\theta_s\right) - \kappa\right) \tag{A4}$$

which is 1 when a given weight $W_{li}$ correctly classifies all the patterns (we will call this a *solution*), and 0 otherwise. The volume of the allowed synapses, which in statistical mechanics is known as the *partition function*, is therefore:

$$Z = \int d\mu(W)\,X_{\xi,\sigma}(W) \tag{A5}$$

where $d\mu(W)$ is the measure over the weights. We will consider in the following

$$\int d\mu(W) \bullet \equiv \int_0^\infty \prod_{li} dW_{li} \bullet \tag{A6}$$

without giving constraints to the norm of the weights. As we will see the norm will be imposed self-consistently by the learning problem.

#### 3. Replica method

To compute the average entropy $\langle \ln Z\rangle_{\xi,\sigma}$ of synaptic weights solutions in the large $N$ limit, we resort the Replica Method [3] that is based on the following identity:

$$\langle \ln Z\rangle_{\xi,\sigma} = \lim_{n\to 0}\frac{\langle Z^n\rangle_{\xi,\sigma} - 1}{n} = \lim_{n\to 0}\frac{1}{n}\ln\langle Z^n\rangle_{\xi,\sigma}$$

This trick reconducts the problem of estimating the log of the partition function in (A5) to the computation of the average of $n$ independent copies of the systems with the same realization of the disorder of the activity patterns and labels $\boldsymbol{\xi}^\mu, \sigma^\mu$:

$$\langle Z^n\rangle = \left\langle \int \prod_{a=1}^{n} d\mu(W^a) \prod_{a,\mu}\theta\left(\frac{\sigma^\mu}{\sqrt{K}}\left(\sum_{l=1}^{K} c_l\, g\left(\sqrt{\frac{K}{N}}\sum_{i=1}^{N/K} W^a_{li}\xi^\mu_{li} - \sqrt{\frac{N}{K}}\theta_d\right) - K\theta_s\right) - \kappa\right)\right\rangle_{\xi,\sigma} . \tag{A7}$$

We will denote from now on $a$ and $b$ as the index that run over replicas $a\,,b=1,\ldots,n$. Notice also that, because of (A4) we can safely consider having labels $\sigma^{\mu}=\pm1$ with the same output coding level as before. The computation follows standard steps [1, 4], which we will sketch here. Firstly we need to perform the average over the activity patterns $\boldsymbol{\xi}^{\mu}$; we can do that by introducing the auxiliary variables

$$\lambda_l^{\mu a}=\sqrt{\frac{K}{N}}\sum_{i=1}^{N/K}W_{li}\xi_{li}^{\mu}-\sqrt{\frac{N}{K}}\theta_d \tag{A8}$$

and the corresponding conjugated variables $\hat{\lambda}_l^{\mu a}$ that arise when we insert the integral representation of the Dirac delta function. The replicated partition function is

$$\begin{aligned}\langle Z^n\rangle=\mathbb{E}_{\sigma}\int\prod_{a}d\mu(W^a)\int\prod_{\mu al}\frac{d\lambda_l^{\mu a}d\hat{\lambda}_l^{\mu a}}{2\pi}e^{i\lambda_l^{\mu a}\hat{\lambda}_l^{\mu a}}\prod_{a,\mu}\theta\left(\frac{\sigma^{\mu}}{\sqrt{K}}\left(\sum_{l=1}^{K}c_l\,g\left(\lambda_l^{\mu a}\right)-K\theta_s\right)-\kappa\right)e^{i\sqrt{\frac{N}{K}}\theta_d\sum_{\mu al}\lambda_l^{\mu a}}.\\ \times\prod_{li\mu}\left\langle e^{-i\xi_{li}^{\mu}\sqrt{\frac{K}{N}}\sum_a W_{li}^a\hat{\lambda}_l^{\mu a}}\right\rangle_{\xi_{li}^{\mu}}.\end{aligned} \tag{A9}$$

The average over patterns can now be performed. In the large $N$ limit, we can use the central limit theorem, having

$$\begin{aligned}\prod_{li\mu}\left\langle e^{-i\xi_{li}^{\mu}\sqrt{\frac{K}{N}}\sum_a W_{li}^a\hat{\lambda}_l^{\mu a}}\right\rangle_{\xi_{li}^{\mu}}&=\prod_{li\mu}\left[1-f_{\text{in}}+f_{\text{in}}e^{-i\sqrt{\frac{K}{N}}\sum_a W_{li}^a\hat{\lambda}_l^{\mu a}}\right]\simeq\\ &=\prod_{li\mu}e^{-if_{\text{in}}\sqrt{\frac{K}{N}}\sum_a W_{li}^a\hat{\lambda}_l^{\mu a}-\frac{f_{\text{in}}(1-f_{\text{in}})K}{2N}\left(\sum_a W_{li}^a\hat{\lambda}_l^{\mu a}\right)^2}\\ &=e^{-if_{\text{in}}\sqrt{\frac{K}{N}}\sum_{\mu al}\hat{\lambda}_l^{\mu a}\sum_i W_{li}^a-\frac{f_{\text{in}}(1-f_{\text{in}})K}{N}\sum_{\mu l}\sum_{a<b}\left(\sum_i W_{li}^aW_{li}^b\right)\hat{\lambda}_l^{\mu a}\hat{\lambda}_l^{\mu b}}\\ &\times e^{-\frac{f_{\text{in}}(1-f_{\text{in}})K}{2N}\sum_{\mu al}\sum_i\left(W_{li}^a\hat{\lambda}_l^{\mu a}\right)^2}\end{aligned} \tag{A10}$$

By defining appropriate order parameters, it is possible to conveniently study the problem in the large-$N$ limit. We define:

$$\sum_i W_{li}^a=\frac{N}{K}\overline{W}+\sqrt{\frac{N}{K}}M_l^a \tag{A11a}$$

$$q_l^{ab}\equiv\frac{K}{N}\sum_i W_{li}^aW_{li}^b \tag{A11b}$$

$$Q_l^a\equiv\frac{K}{N}\sum_i\left(W_{li}^a\right)^2 \tag{A11c}$$

(A11a) represents the average synaptic weight; we expressed it in two contributions. The first one represents an averaged *scaled* synaptic weight

$$\overline{W}=\frac{\theta_d}{f_{\text{in}}} \tag{A12}$$

justified by the fact that for each sub-perceptron of the first layer only $\frac{N}{K}f_{\text{in}}$ synapses contribute. The second term instead is a $\sqrt{\frac{N}{K}}$ correction that is needed in order to fine tune the average synaptic weight relatively to the threshold $\theta_d$.

The quantity $q_l^{ab}$ is the overlap between the weights of two different replicas $a$ and $b$ belonging to the same dendritic branch $l$, while $Q_l^a$ represents the averaged squared norm of a synaptic weight belonging to dendritic branch $l$.

Enforcing the definitions (A11) in (A9), by using Dirac delta functions, we can express the replicated partition function as an integration over the order parameters $q_l^{ab},\hat{q},Q,\hat{Q},M,\hat{M}$:

$$\begin{aligned}\langle Z^n\rangle_{\xi,\sigma}=\int\prod_{a<b,l}\frac{dq_l^{ab}d\hat{q}_l^{ab}}{2\pi K/N}\int\prod_{a,l}\frac{dQ_l^ad\hat{Q}_l^a}{2\pi K/N}\int\prod_{a,l}\frac{dM_l^ad\hat{M}_l^a}{2\pi\sqrt{K/N}}e^{-\frac{N}{K}\sum\limits_{a<b,l}q_l^{ab}\hat{q}_l^{ab}-\frac{N}{K}\sum\limits_{a,l}Q_l^a\hat{Q}_l^a-\frac{N}{K}\overline{W}\sum\limits_{a,l}\hat{M}_l^a}\\ \times e^{\frac{N}{K}G_S\left(\hat{q}_l^{ab},\hat{Q}_l^a,\hat{M}_l^a\right)+N\alpha G_E\left(q_l^{ab},Q_l^a,M_l^a\right)}.\end{aligned} \tag{A13}$$

where we collected the entropic contribution $G_S$ and the energetic one $G_E$. The first is the usual term that counts how many coupling vectors $W^a$ fulfill the constraints (A11); the second is specific to the learning rule which is used, and depends on the Heaviside function that counts learned patterns:

$$G_S\left(\hat{q}_l^{ab}, \hat{Q}_l^a, \hat{M}_l^a\right) = \ln \int_0^\infty \prod_{a,l} dW_l^a \, e^{\sum\limits_{a<b,l} \hat{q}_l^{ab} W_l^a W_l^b + \sum\limits_{a,l} \hat{Q}_l^a (W_l^a)^2 + \sum\limits_{a,l} \hat{M}_l^a W_l^a} \tag{A14a}$$

$$G_E\left(q_l^{ab}, Q_l^a, M_l^a\right) = \ln \mathbb{E}_\sigma \int \prod_{a,l} \frac{d\lambda_l^a d\hat{\lambda}_l^a}{2\pi} \Theta\left(\frac{\sigma}{\sqrt{K}}\left(\sum_l c_l \, g(\lambda_l^a) - K\theta_s\right) - \kappa\right) e^{i\sum\limits_{a,l} \lambda_l^a \hat{\lambda}_l^a - f_{\text{in}}(1-f_{\text{in}}) \sum\limits_{a<b,l} q_l^{ab} \hat{\lambda}_l^a \hat{\lambda}_l^b} \tag{A14b}$$

$$\times \, e^{-\frac{f_{\text{in}}(1-f_{\text{in}})}{2} \sum\limits_{a,l} Q_l^a (\hat{\lambda}_l^a)^2 - i f_{\text{in}} \sum\limits_{a,l} M_l^a \hat{\lambda}_l^a} .$$

We can now evaluate in the large $N$ limit (A13) using the saddle point method. In order restrict the space where to search saddle points we proceed by assuming a particular form of the order parameters, which is the main topic of the next section.

#### 4. Replica Symmetric analysis

We use a Replica Symmetric (RS) ansatz, i.e. we assume that the order parameters do not depend on the replica indexes and of the index corresponding to the dendritic branch:

$$q_l^{ab} = q \,, \qquad Q_l^a = Q \,, \qquad M_l^a = M \,, \tag{A15a}$$

$$\hat{q}_l^{ab} = \hat{q} \,, \qquad \hat{Q}_l^a = \hat{Q} \,, \qquad \hat{M}_l^a = \hat{M} \tag{A15b}$$

In the RS ansatz and in the small $n$ limit, using *Hubbard-Stratonovich* transformations:

$$e^{\frac{1}{2}bx^2} = \int \frac{dz}{\sqrt{2\pi}} \, e^{-\frac{z^2}{2} + \sqrt{b}xz} \,,$$

the entropic and energetic terms are the following

$$\mathscr{G}_S \equiv \lim_{n\to 0} \frac{1}{nK} G_S(\hat{q}, \hat{Q}, \hat{M}) = \ln \sqrt{\frac{2\pi}{\hat{q} - 2\hat{Q}}} + \frac{1}{2}\left(\frac{\hat{M}^2 + \hat{q}}{\hat{q} - 2\hat{Q}}\right) + \int Dz \, \ln H\left(-\frac{\hat{M} + \sqrt{\hat{q}}z}{\sqrt{\hat{q} - 2\hat{Q}}}\right) \tag{A16a}$$

$$\mathscr{G}_E \equiv \lim_{n\to 0} \frac{G_E(q, Q, M)}{n} = \mathop{\mathbb{E}}_\sigma \int \prod_l Dt_l \, \ln\left[\int \prod_l D\lambda_l \, \Theta\left(\frac{\sigma}{\sqrt{K}}\left(\sum_l c_l \, g\left(\sqrt{f_{\text{in}}(1-f_{\text{in}})(Q-q)}\lambda_l + a_l\right) - K\theta_s\right) - \kappa\right)\right] \tag{A16b}$$

where we have introduced the variable

$$a_l \equiv f_{\text{in}} M + \sqrt{f_{\text{in}}(1-f_{\text{in}})q}\, t_l \tag{A17}$$

for convenience. In (A16a) we have also introduced $H(x) \equiv \int_x^\infty Dz = \frac{1}{2}\text{Erfc}\left(\frac{x}{\sqrt{2}}\right)$ and $Dz \equiv G(z)\,dz$ with $G(z)$ being a standard normal Gaussian $G(z) = \exp(-z^2/2)/\sqrt{2\pi}$.

##### *a. Large K limit*

We focus on the limit $K \to \infty$ $\left(\text{with } \frac{K}{N} \to 0\right)$, for two main reasons:

- on the analytical level, it allows to simplify the numerical evaluation of the saddle point equations corresponding to the RS ansatz;

- it is *biologically* realistic: the number of dendritic branches in neurons is typically large (in some cases even more than a hundred, REF) and the number of synapses in each branch is typically large as well (REFS?).

To evaluate this limit, we need to do some manipulations on the energetic contribution (A16b) which are based on the central limit theorem. Let us first consider the term in square brackets:

$$\begin{aligned} I &= \int \prod_l D\lambda_l\, \Theta\left(\frac{\sigma}{\sqrt{K}}\left(\sum_l c_l\, g\left(\sqrt{f_{\rm in}(1-f_{\rm in})(Q-q)}\lambda_l + a_l\right) - K\theta_s\right) - \kappa\right) \\ &= \int \frac{dh\, d\hat{h}}{2\pi} e^{-ih\hat{h}}\, \Theta\left[\sigma\left(h - \sqrt{K}\theta_s\right) - \kappa\right] \int \prod_l D\lambda_l\, e^{\frac{i\hat{h}}{\sqrt{K}}\sum_l c_l\, g\left(a_l + \sqrt{f_{\rm in}(1-f_{\rm in})(Q-q)}\,\lambda_l\right)} \end{aligned} \tag{A18}$$

In the large $K$ limit we can therefore expand the exponential up to second order

$$\begin{aligned} I \simeq \int \frac{dh\, d\hat{h}}{2\pi} e^{-ih\hat{h}}\, \Theta\left(\sigma(h - \sqrt{K}\theta_s) - \kappa\right) \int \prod_l D\lambda_l \Bigg[ 1 + \frac{i\hat{h}}{\sqrt{K}} \sum_l c_l\, g\left(a_l + \sqrt{f_{\rm in}(1-f_{\rm in})(Q-q)}\,\lambda_l\right) + \\ - \frac{\hat{h}^2}{2K}\left(\sum_l c_l\, g\left(a_l + \sqrt{f_{\rm in}(1-f_{\rm in})(Q-q)}\,\lambda_l\right)\right)^2 \Bigg]. \end{aligned} \tag{A19}$$

and we can integrate with respect to all the $\lambda_l$ variables term by term. Exponentiating the expression again and integrating over $\hat{h}$ we get

$$I = \int Dh\, \Theta\left(\sigma(M^{(0)} + \sqrt{\Delta^{(0)}} h - \sqrt{K}\theta_s) - \kappa\right) \tag{A20}$$

where we have introduced the variables:

$$M^{(0)} = \frac{1}{\sqrt{K}} \sum_l c_l \langle g \rangle_\lambda \tag{A21a}$$

$$D^{(0)} = \frac{1}{K} \sum_l c_l^2 \left[ \langle g^2 \rangle_\lambda - \langle g \rangle_\lambda^2 \right]. \tag{A21b}$$

and the notation

$$\langle g \rangle_\lambda = \int D\lambda\, g\left(f_{\rm in} M + \sqrt{f_{\rm in}(1-f_{\rm in})q}\, t_l + \sqrt{f_{\rm in}(1-f_{\rm in})(Q-q)}\,\lambda\right). \tag{A22}$$

Using again the central limit theorem for the $K$ integrals over the variable $t_l$ we get:

$$\mathcal{G}_E = \mathop{\mathbb{E}}_{\sigma} \int Dt \ln \int D\lambda\, \Theta\left[\sigma\left(M_0 + \sqrt{D_0}\, t + \sqrt{D_1}\,\lambda - \sqrt{K}\theta_s\right) - \kappa\right] \tag{A23}$$

where:

$$M_0 = \frac{1}{\sqrt{K}} \sum_l c_l \langle \langle g \rangle_\lambda \rangle_t = m_c \langle \langle g \rangle_\lambda \rangle_t \tag{A24a}$$

$$D_0 = \frac{1}{K} \sum_l c_l^2 \left[ \langle \langle g \rangle_\lambda^2 \rangle_t - \langle \langle g \rangle_\lambda \rangle_t^2 \right] = w_c \left[ \langle \langle g \rangle_\lambda^2 \rangle_t - \langle \langle g \rangle_\lambda \rangle_t^2 \right] \tag{A24b}$$

$$D_1 = w_c \left[ \langle \langle g^2 \rangle_\lambda \rangle_t - \langle \langle g \rangle_\lambda^2 \rangle_t \right] \tag{A24c}$$

and for example

$$\langle \langle g \rangle_\lambda \rangle_t = \int Dt D\lambda\, g\left(f_{\rm in} M + \sqrt{f_{\rm in}(1-f_{\rm in})q}\, t + \sqrt{f_{\rm in}(1-f_{\rm in})(Q-q)}\,\lambda\right). \tag{A25}$$

Using the definition of the committee machine in which all weights in the second layer are s.t. $c_l = 1$, we have $m_c = \sqrt{K}$ and $w_c = 1$. In order to have a well-defined large $K$ limit we have to impose (analogously to what we have done on the dendritic threshold $\theta_d$) that the divergence induced by the somatic threshold $\theta_s$ cancels with the one coming from $M_0$. We therefore impose that $M$ scales, in the large $K$ limit, as

$$M = \overline{M} + \frac{\delta M}{\sqrt{K}}\,. \tag{A26}$$

$M_0$ can be simplified by making a rotation over the integration measures $\lambda$ and $t$

$$\begin{aligned} M_0 = \sqrt{K}\langle\langle g\rangle_\lambda\rangle_t &= \sqrt{K}\int D\lambda\, Dt\, g\left(\sqrt{f_{\rm in}(1-f_{\rm in})q}\,t + \sqrt{f_{\rm in}(1-f_{\rm in})(Q-q)}\lambda + fM\right) \\ &= \sqrt{K}\int D\lambda\, g\left(\sqrt{f_{\rm in}(1-f_{\rm in})Q}\,\lambda + f_{\rm in}M\right) \end{aligned} \tag{A27}$$

We can now insert the scaling in (A26)

$$\begin{aligned} M_0 &= \sqrt{K}\int D\lambda\, g\left(\sqrt{f_{\rm in}(1-f_{\rm in})Q}\,\lambda + f_{\rm in}M\right) \\ &\simeq \sqrt{K}\int D\lambda\, g\left(\sqrt{f_{\rm in}(1-f_{\rm in})Q}\lambda + f_{\rm in}\overline{M}\right) + f_{\rm in}\,\delta M\int D\lambda\, g'\left(\sqrt{f_{\rm in}(1-f_{\rm in})Q}\,\lambda + f_{\rm in}\overline{M}\right) \\ &= \sqrt{K}\,\theta_s + \Delta \end{aligned} \tag{A28}$$

Therefore, $\overline{M}$ is fixed by the relation:

$$\theta_s = \int Dy\, g\left(\sqrt{f_{\rm in}(1-f_{\rm in})Q}\,y + f_{\rm in}\overline{M}\right)\,, \tag{A29}$$

that involves the output threshold. Hence the energetic term (A23) becomes:

$$\mathcal{G}_E = \mathop{\mathbb{E}}_{\sigma}\int Dz\,\ln H\left(\frac{\kappa-\sigma\Delta+\sqrt{D_0}\,z}{\sqrt{D_1}}\right) = \int Dz\,\left[f_{\rm out}\ln H\left(\frac{\kappa-\Delta+\sqrt{D_0}\,z}{\sqrt{D_1}}\right) + (1-f_{\rm out})\ln H\left(\frac{\kappa+\Delta+\sqrt{D_0}\,z}{\sqrt{D_1}}\right)\right] \tag{A30}$$

where the order parameters, strictly dependent on the choice of the activation function $g(x)$, are simplified as:

$$\begin{aligned} \Delta &\equiv f_{\rm in}M\int Dx\, g'\left(\sqrt{f_{\rm in}(1-f_{\rm in})Q}\,x + f_{\rm in}\overline{M}\right) \\ D_0 &= \Delta_q - \Delta_0 \\ D_1 &= \Delta_Q - \Delta_q \end{aligned}$$

where we have renamed $\delta M$ by $M$ with a slight abuse of notation. We have also defined the generic "effective order parameter" or *kernel* functions

$$\Delta_q = \int Dx\left[\int Dy\, g\left(\sqrt{f_{\rm in}(1-f_{\rm in})q}\,x + \sqrt{f_{\rm in}(1-f_{\rm in})(Q-q)}\,y + f_{\rm in}\overline{M}\right)\right]^2 \tag{A32}$$

$\Delta_Q$ and $\Delta_0$ being obtained by simply substituting in the previous expression $q \to Q$ and $q \to 0$ respectively. We report them here for clarity

$$\Delta_Q = \int Dx\, g^2\left(\sqrt{f_{\rm in}(1-f_{\rm in})Q}\,x + f_{\rm in}\overline{M}\right)\,, \tag{A33a}$$

$$\Delta_0 = \left[\int Dy\, g\left(\sqrt{f_{\rm in}(1-f_{\rm in})Q}\,y + f_{\rm in}\overline{M}\right)\right]^2\,. \tag{A33b}$$

We have called $\Delta_q$ an *effective order parameter* since (A30) (and therefore the whole quenched entropy) is perfectly equivalent to the one found in the perceptron model studied by Brunel in a series of papers [5, 6], but where order parameters $q$ and $Q$ are substituted respectively by $\Delta_q - \Delta_0$ and $\Delta_Q - \Delta_0$.

#### b. Free entropy and saddle point equations

The average *free entropy* of the dendritic model of a neuron is therefore

$$\phi = \lim_{N\to\infty} \frac{1}{N}\langle \ln Z\rangle_{\xi,\sigma} = \frac{q\hat{q}}{2} - Q\hat{Q} - \overline{W}\hat{M} + \mathcal{G}_S + \alpha\,\mathcal{G}_E \tag{A34}$$

We now need to compute the saddle point equations by differentiating (A34) with respect to the order parameters $Q, q, M, \hat{Q}, \hat{q}, \hat{M}$. The saddle point equations involving the entropic term (i.e. taking derivatives with respect to $\hat{M}$, $\hat{Q}$ and $\hat{q}$) are the same as in the case of the perceptron

$$\overline{W} = \int Dz\, \frac{\int_0^\infty dW\, W e^{-(\hat{q}-2\hat{Q})\frac{W^2}{2}+(\hat{M}+\sqrt{\hat{q}}z)W}}{\int_0^\infty dW\, e^{-(\hat{q}-2\hat{Q})\frac{W^2}{2}+(\hat{M}+\sqrt{\hat{q}}z)W}} \tag{A35a}$$

$$Q = \int Dz\, \frac{\int_0^\infty dW\, W^2 e^{-(\hat{q}-2\hat{Q})\frac{W^2}{2}+(\hat{M}+\sqrt{\hat{q}}z)W}}{\int_0^\infty dW\, e^{-(\hat{q}-2\hat{Q})\frac{W^2}{2}+(\hat{M}+\sqrt{\hat{q}}z)W}} \tag{A35b}$$

$$q = \int Dz\, \frac{\int_0^\infty dW\, \left(W^2 - \frac{zW}{\sqrt{\hat{q}}}\right) e^{-(\hat{q}-2\hat{Q})\frac{W^2}{2}+(\hat{M}+\sqrt{\hat{q}}z)W}}{\int_0^\infty dW\, e^{-(\hat{q}-2\hat{Q})\frac{W^2}{2}+(\hat{M}+\sqrt{\hat{q}}z)W}} = \int Dz \left[\frac{\int_0^\infty dW\, W e^{-(\hat{q}-2\hat{Q})\frac{W^2}{2}+(\hat{M}+\sqrt{\hat{q}}z)W}}{\int_0^\infty dW\, e^{-(\hat{q}-2\hat{Q})\frac{W^2}{2}+(\hat{M}+\sqrt{\hat{q}}z)W}}\right]^2 \tag{A35c}$$

In order to express the remaining saddle point equations in a compact way, we define the quantities:

$$a_\sigma(z) = \frac{\sqrt{D_0}z - \sigma\Delta + \kappa}{\sqrt{D_1}} = \sqrt{\frac{D_0}{D_1}}(z - \tau_\sigma)$$
$$\tau_\sigma = \frac{\sigma\Delta - \kappa}{\sqrt{D_0}}\,.$$

Deriving (A34) with respect to $M$, $Q$ and $q$ lead respectively to

$$0 = \mathbb{E}_\sigma \sigma \int Dz\, \frac{G(a_\sigma(z))}{H(a_\sigma(z))} \tag{A37a}$$

$$\hat{Q} = \frac{\alpha}{2}\mathbb{E}_\sigma \int Dz\, \frac{G(a_\sigma(z))}{H(a_\sigma(z))}\left[\frac{a_\sigma(z)}{D_1}\frac{dD_1}{dQ} - \frac{z}{\sqrt{D_0 D_1}}\frac{dD_0}{dQ}\right] \tag{A37b}$$

$$\hat{q} = \alpha\mathbb{E}_\sigma \int Dz\, \frac{G(a_\sigma(z))}{H(a_\sigma(z))}\left[-\frac{a_\sigma(z)}{D_1}\frac{dD_1}{dq} + \frac{z}{\sqrt{D_0 D_1}}\frac{dD_0}{dq}\right] \tag{A37c}$$

where we used the saddle point equation (A37a) when performing the derivative in $Q$[1]. The six saddle points Eqs. ((A35a), A35b, A35c, A37a, A37b, A37c), obtained in the large $K, N \to \infty$ limit with $N \gg K$, have to be numerically solved to obtain the values of the order parameters and represent the final result of our RS analysis.

Notice that imposing $g(x) = x$ we recover the previous saddle point expressions obtained for the simple linear neuron model [5, 6]. Notice also that in the case $f_{\text{out}} = \frac{1}{2}$ saddle point equation (A37a) gives $M = 0$; in this case therefore $\tau_\sigma = 0$.

#### c. Effective order parameters for some non-linearities

We report here the analytical expressions of the effective order parameters for several non-linearities of interest

- *Recovering the one-layer neuron model*: if we impose $g(x) = x$ we recover the one-layer neuron model. We report here the expressions of the corresponding effective order parameters for convenience

$$\Delta = f_{\text{in}} M \tag{A38a}$$

[1] this is why no derivative with respect to $Q$ of $\Delta$ compares in (A37b)

$$\Delta_q = f_{\rm in}(1-f_{\rm in})q + f_{\rm in}^2\overline{M}^2 \tag{A38b}$$

and, in particular

$$\Delta_Q = f_{\rm in}(1-f_{\rm in})Q + f_{\rm in}^2\overline{M}^2 \tag{A39a}$$
$$\Delta_0 = f_{\rm in}^2\overline{M}^2 \tag{A39b}$$

As a result $\overline{M}$ does not appear anywhere in the energetic term of equation (A30), and $\theta_s = f_{\rm in}\overline{M}$ is also irrelevant.

- *Theta non-linearity*: $g(x) = \Theta(x)$.

  To evaluate the integrals it is useful to use the following identity

$$\int Dz\, H^2\left(a+bz\right) = H\left(\frac{a}{\sqrt{1+b^2}}\right) - 2T\left(\frac{a}{\sqrt{1+b^2}}, \frac{1}{\sqrt{1+2b^2}}\right) \tag{A40}$$

  where $T$ is the Owen's $T$ function defined as

$$T(h,s) \equiv \frac{1}{2\pi}\int_0^s dx\, \frac{e^{-(1+x^2)\frac{h^2}{2}}}{1+x^2}\,, \tag{A41}$$

  and has the following important properties

$$T(h,s) \simeq \frac{G(h)s}{\sqrt{2\pi}} + O\left(s^2\right) \qquad \text{for } s\to 0 \tag{A42a}$$
$$T(h,1) = \frac{1}{2}H(h)H(-h) \tag{A42b}$$

  where we remind that $G(x)$ is the Gaussian with mean zero and unit variance. Defining the quantity

$$M_\star = \frac{f_{\rm in}\overline{M}}{\sqrt{f_{\rm in}(1-f_{\rm in})Q}} \tag{A43}$$

  the effective order parameters for the theta non-linearity are

$$\Delta = M_\star G\left(-M_\star\right) \tag{A44a}$$
$$\Delta_q = H(-M_\star) - 2T\left(M_\star, \sqrt{\frac{Q-q}{Q+q}}\right) \tag{A44b}$$

  and, in particular

$$\Delta_Q = H(-M_\star)\,, \tag{A45a}$$
$$\Delta_0 = H(-M_\star)^2\,. \tag{A45b}$$

  $\overline{M}$ is fixed by the relation

$$\theta_s = H(-M_\star)\,. \tag{A46}$$

- *ReLU non-linearity*: $g(x) = x\Theta(x)$ We have

$$\Delta = f_{\rm in}MH\left(-M_\star\right) \tag{A47a}$$
$$\Delta_q = f_{\rm in}(1-f_{\rm in})(Q-q)\sqrt{\frac{Q-q}{Q+q}}G^2\left(\frac{f_{\rm in}\overline{M}}{\sqrt{f_{\rm in}(1-f_{\rm in})(Q+q)}}\right) + f_{\rm in}\left((1-f_{\rm in})q + f_{\rm in}\overline{M}^2\right)\left[H\left(-M_\star\right) - 2T\left(-M_\star, \sqrt{\frac{Q-q}{Q+q}}\right)\right] \tag{A47b}$$

$$+ 2f_{\rm in}\overline{M}\sqrt{f_{\rm in}(1-f_{\rm in})Q}\,G\left(M_\star\right)H\left(-M_\star\sqrt{\frac{Q-q}{Q+q}}\right) + 2f_{\rm in}(1-f_{\rm in})q\sqrt{\frac{Q-q}{Q+q}}G\left(M_\star\right)G\left(M_\star\sqrt{\frac{Q-q}{Q+q}}\right)$$

and in particular

$$\Delta_Q = f_{\rm in}\left((1-f_{\rm in})Q + f_{\rm in}\overline{M}^2\right)H\left(-M_\star\right) + f_{\rm in}\overline{M}\sqrt{f_{\rm in}(1-f_{\rm in})Q}\,G\left(-M_\star\right)\,, \tag{A48a}$$

$$\Delta_0 = \left[\sqrt{f_{\rm in}(1-f_{\rm in})Q}\,G\left(M_\star\right) + f_{\rm in}\overline{M}H\left(-M_\star\right)\right]^2 \tag{A48b}$$

Note that $\overline{M}$ is fixed by the relation

$$\theta_s = \sqrt{f_{\rm in}(1-f_{\rm in})Q}\,G\left(M_\star\right) + f_{\rm in}\overline{M}H\left(-M_\star\right) \tag{A49}$$

#### *d. Small data regime*

In the $\alpha \to 0$ limit the saddle point equations can be solved exactly. Indeed $\hat{q} = \hat{Q} = 0$ and equations (A35a), (A35b), (A35c) give respectively

$$\overline{W} = \frac{\theta_d}{f_{\rm in}} = \frac{\int_0^\infty dW\,W e^{\hat{M}W}}{\int_0^\infty dW\,e^{\hat{M}W}} = -\frac{1}{\hat{M}} \tag{A50a}$$

$$Q = \frac{\int_0^\infty dW\,W^2 e^{\hat{M}W}}{\int_0^\infty dW\,e^{\hat{M}W}} = \frac{2}{\hat{M}^2} \tag{A50b}$$

$$q = \left[\frac{\int_0^\infty dW\,W e^{\hat{M}W}}{\int_0^\infty dW\,e^{\hat{M}W}}\right]^2 = \frac{1}{\hat{M}^2} \tag{A50c}$$

#### *e. Distribution of dendritic preactivations*

We derive here the distribution of dendritic preactivations after learning. Since the each dendritic branch has access to an indepedent portion of the input, the distribution of the preactivations if factorized over the $K$ dendritic branches. In the large $K$ limit the distribution tends to a Gaussian $\mathcal{N}(\mu,\sigma)$ as can be inspected from the post activation mean (A29) and from the argument of the kernel functions (A32). Denoting by $\lambda_l \equiv \sqrt{\frac{K}{N}}\sum_{i=1}^{N/K} W_{li}\xi_{li} - \sqrt{\frac{N}{K}}\theta_d$ as in the main text, the mean and the variance of the distribution of the $l-$th dendritic branch are respectively

$$\mu = \mathbb{E}_{\boldsymbol{\xi}}\lambda_l = f_{\rm in}\overline{M} \tag{A51a}$$

$$\sigma^2 = \mathbb{E}_{\boldsymbol{\xi}}\lambda_l^2 - \mu^2 = f_{\rm in}(1-f_{\rm in})Q \tag{A51b}$$

as confirmed by (A29).

In the $\alpha \to 0$ the variance above can be expressed explicitly in terms of $\theta_d$ and $f_{\rm in}$ thanks to (A50a) and (A50b). We get that the standard deviation of the dendritic preactivation depends linearly on the dendritic inhibition threshold

$$\sigma = \theta_d\sqrt{\frac{2(1-f_{\rm in})}{f_{\rm in}}}\,. \tag{A52}$$

We checked that this linear relation is satisfied even at finite $\alpha$, or considering a different distribution over the weights at initialization, see below. This shows that if $\theta_d$ is small one does not completely use the non-linearity and the model behaves like a one-layer model.

#### *f. Limit of large somatic thresholds*

When the somatic threshold $\theta_s$ diverges, the only way to satisfy (A29) is that $\overline{M}$ diverges if the non-linearity $g$ is unbounded. This can be inspected already in (A49) in the case of the ReLU non-linearity.

Instead if the non-linearity is bounded, the right hand side of (A29) is a bounded function of $\overline{M}$ as well, therefore above a certain critical value of $\theta_s$ it will not possible to find the corresponding value of $\overline{M}$.

Moreover, if the non-linearity diverges linearly for large arguments, we expect to recover back the free energy and the expressions for the one-layer neuron model for large $\theta_s$. Indeed expanding equation (A29) one finds $\overline{M} \sim \theta_s / f_{\rm in}$ and the effective order parameters reduce to those ones of the perceptron, see (A38).

#### 5. Critical capacity

In this section we show how in our formalism, it is possible to compute the maximal number of inputs that the neuron is able to classify. We underline that this can be done for a generic form of dendritic non-linearity.

In the critical capacity limit, the set of possible synaptic weights shrinks towards a single point and $q$ tends to $Q$:

$$q = Q - dq. \tag{A53}$$

Correspondingly, the other order parameters scale as

$$\hat{q}\,, \hat{Q} \sim \frac{C}{dq^2} \tag{A54a}$$

$$\hat{q} - 2\hat{Q} \sim \frac{A}{dq} \tag{A54b}$$

$$\hat{M} \sim -\frac{B\sqrt{C}}{dq} \tag{A54c}$$

Using the identities (where $a$ is a positive constant)

$$\frac{\int_0^\infty dx\, x\, e^{-a\frac{x^2}{2}+bx}}{\int_0^\infty dx\, e^{-a\frac{x^2}{2}+bx}} = \frac{b}{a} + \frac{1}{\sqrt{a}} \frac{G\left(-\frac{b}{\sqrt{a}}\right)}{H\left(-\frac{b}{\sqrt{a}}\right)} \tag{A55a}$$

$$\frac{\int_0^\infty dx\, x^2\, e^{-a\frac{x^2}{2}+bx}}{\int_0^\infty dx\, e^{-a\frac{x^2}{2}+bx}} = \frac{1}{a} + \frac{b^2}{a^2} + \frac{b}{a^{3/2}} \frac{G\left(-\frac{b}{\sqrt{a}}\right)}{H\left(-\frac{b}{\sqrt{a}}\right)} \tag{A55b}$$

and the expansion

$$\frac{G(x)}{H(x)} \simeq x\theta(x)\,, \qquad \text{for } |x| \gg 1\,, \tag{A56}$$

the saddle point equations (A35) can be written as

$$\overline{W} = \frac{\sqrt{C}}{A}\left[G(B) - BH(B)\right] \tag{A57a}$$

$$Q = \frac{C}{A^2}\left[\left(1+B^2\right)H(B) - BG(B)\right] \tag{A57b}$$

$$A = H(B). \tag{A57c}$$

More work is required to derive the asymptotic limit of equations (A37). First of all we need the expansion of the effective order parameters

$$D_0 = \Delta_q - \Delta_0 = \Delta_Q - \Delta_0 - D_1 \simeq \Gamma_0 - \Gamma_1 dq \tag{A58a}$$

$$D_1 = \Delta_Q - \Delta_q = \Gamma_1 dq + O(dq^2) \tag{A58b}$$

where we have defined

$$\Gamma_0 = \Delta_Q - \Delta_0 = \int Dx\, g^2\left(\sqrt{f_{\rm in}(1-f_{\rm in})Q}\, x + f_{\rm in}\overline{M}\right) - \left[\int Dy\, g\left(\sqrt{f_{\rm in}(1-f_{\rm in})Q}\, y + f_{\rm in}\overline{M}\right)\right]^2 \tag{A59a}$$

$$\Gamma_1 = f_{\rm in}(1-f_{\rm in}) \int Dz\, \left[g'\left(\sqrt{f_{\rm in}(1-f_{\rm in})Q}\, z + f_{\rm in}\overline{M}\right)\right]^2 \tag{A59b}$$

Now we subtract (A37b) with (A37c) getting

$$\begin{aligned}\hat{q}-2\hat{Q} &= \alpha\mathbb{E}_\sigma \int Dz\, \frac{G(a_\sigma(z))}{H(a_\sigma(z))}\left[\frac{z}{\sqrt{D_0D_1}}\left(\frac{dD_0}{dq}+\frac{dD_0}{dQ}\right)-\frac{a_\sigma(z)}{D_1}\left(\frac{dD_1}{dq}+\frac{dD_1}{dQ}\right)\right] \\ &= \alpha\mathbb{E}_\sigma \int Dz\, \frac{G(a_\sigma(z))}{H(a_\sigma(z))}\left[\frac{z}{\sqrt{D_0D_1}}\frac{d\Gamma_0}{dQ}-\frac{a_\sigma(z)}{D_1}\frac{d\Gamma_1}{dQ}dq\right]\end{aligned} \tag{A60}$$

Similarly (A37c) becomes

$$\hat{q} = \alpha\mathbb{E}_\sigma \int Dz\, \frac{G(a_\sigma(z))}{H(a_\sigma(z))}\left[-\frac{a_\sigma(z)}{D_1}\frac{dD_1}{dq}\right] \tag{A61}$$

because the second term in (A37c) is subleading in $dq$. Using the expansion (A56) and the identities

$$\int Dz\, z^2\,\Theta(z-\tau_\sigma) = H(\tau_\sigma)+\tau_\sigma G(\tau_\sigma) \tag{A62a}$$

$$\int Dz\, z\,\Theta(z-\tau_\sigma) = G(\tau_\sigma) \tag{A62b}$$

we obtain the following saddle point equations

$$0 = \mathbb{E}_\sigma \sigma\left[G(\tau_\sigma)-\tau_\sigma H(\tau_\sigma)\right] \tag{A63a}$$

$$A = \alpha_c\left[\frac{1}{\Gamma_1}\frac{d\Gamma_0}{dQ}\,\mathbb{E}_\sigma H(\tau_\sigma)-\frac{\Gamma_0}{\Gamma_1^2}\frac{d\Gamma_1}{dQ}\mathbb{E}_\sigma\left[(1+\tau_\sigma^2)H(\tau_\sigma)-\tau_\sigma G(\tau_\sigma)\right]\right] \tag{A63b}$$

$$C = \frac{\alpha_c\Gamma_0}{\Gamma_1}\mathbb{E}_\sigma\left[(1+\tau_\sigma^2)H(\tau_\sigma)-\tau_\sigma G(\tau_\sigma)\right]\,, \tag{A63c}$$

which involve the critical capacity as an unknown parameter to find. Notice that in the previous equation we have redefined $\tau_\sigma = (\sigma\Delta-\kappa)/\sqrt{\Gamma_0}$. The full set of saddle point equations for the order parameters $A$, $B$, $C$, $M$, $Q$, $\overline{M}$ and for $\alpha_c$ are

$$\overline{W} = \frac{\sqrt{C}}{A}\left[G(B)-BH(B)\right] \tag{A64a}$$

$$Q = \frac{C}{A^2}\left[\left(1+B^2\right)H(B)-BG(B)\right] \tag{A64b}$$

$$A = H(B) \tag{A64c}$$

$$0 = \mathbb{E}_\sigma\sigma\left[G(\tau_\sigma)-\tau_\sigma H(\tau_\sigma)\right] \tag{A64d}$$

$$A = \frac{\alpha_c}{\Gamma_1}\frac{d\Gamma_0}{dQ}\,\mathbb{E}_\sigma H(\tau_\sigma)-\frac{1}{\Gamma_1}\frac{d\Gamma_1}{dQ}C \tag{A64e}$$

$$C = \frac{\alpha_c\Gamma_0}{\Gamma_1}\mathbb{E}_\sigma\left[(1+\tau_\sigma^2)H(\tau_\sigma)-\tau_\sigma G(\tau_\sigma)\right] \tag{A64f}$$

$$\theta_s = \int Dy\, g\left(\sqrt{f_{\rm in}(1-f_{\rm in})Q}\,y+f_{\rm in}\overline{M}\right)\,. \tag{A64g}$$

As we have anticipated before, in the case $f_{\rm out}=0.5$ the equations can be further simplified, since $M=0$; if also $\kappa=0$ therefore $\tau_\sigma=0$; the saddle point equations (A64) then reduce to

$$\overline{W} = \sqrt{\frac{\alpha_c\Gamma_0}{2\Gamma_1}}\frac{1}{H(B)}\left[G(B)-BH(B)\right] \tag{A65a}$$

$$Q = \frac{\alpha_c\Gamma_0}{2\Gamma_1}\frac{1}{H^2(B)}\left[\left(1+B^2\right)H(B)-BG(B)\right] \tag{A65b}$$

$$\alpha_c = \frac{2\Gamma_1 H(-B)}{\frac{d\Gamma_0}{dQ}-\frac{\Gamma_0}{\Gamma_1}\frac{d\Gamma_1}{dQ}} \tag{A65c}$$

$$\theta_s = \int Dy\, g\left(\sqrt{f_{\rm in}(1-f_{\rm in})Q}\,y+f_{\rm in}\overline{M}\right)\,. \tag{A65d}$$

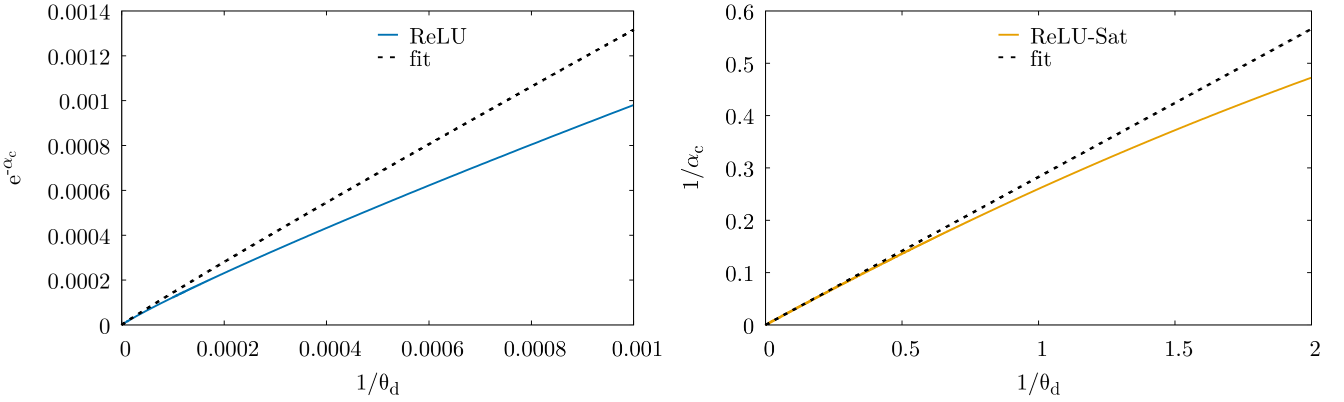


(a) ReLU.

(b) ReLU-Sat.

Figure 1. Fit to the critical capacity for large dendritic thresholds for the ReLU (left panel) and the ReLU-Sat (right panel) non-linearities. The external parameters are the same used in the corresponding figures of the main text, i.e. $f_{\text{in}} = f_{\text{out}} = \theta_s = 0.5$ and $\kappa = 0$. Notice that in the case of the ReLU function (left panel) we are plotting $e^{-\alpha_c}$ versus $\theta_d$ at variance to the ReLU-Sat. In each plot the dashed black line represents a linear fit $a + bx$ of the analytical data for large $\theta_d$.

#### a. *Limit of large dendritic threshold*

As mentioned in the main text, the critical capacity of the model depends strongly on the shape of the non-linearity. In particular, in the limit of large dendritic threshold the critical capacity can diverge differently depending if the non-linearity saturates or not for a sufficiently large stimulus. We show in Fig. 1 how the critical capacity diverges logarithmically in $\theta_d$ for the ReLU activation function whereas the divergence is linear in the ReLU-Sat activation function. Performing a fit we get when $\theta_d \to \infty$

$$\alpha_c^{\text{ReLU-Sat}} \simeq 3.518\,\theta_d \tag{A66a}$$

$$\alpha_c^{\text{ReLU}} \simeq 0.9602\,\ln \theta_d \tag{A66b}$$

#### b. *Limit of small dendritic threshold*

As shown in the main text numerically and above for $\alpha < \alpha_c$, in the low dendritic threshold regime for fixed $\theta_s$ the model with Polsky and ReLU activation function behaves like a one-layer model. We give another quantitative argument here for $\alpha = \alpha_c$. Since $\theta_d \to 0$ the right-hand side of the first of (A64) should go to zero. Since the function $G(B) - BH(B)$ does not go to zero for finite values of $B$, by necessity $C \to 0$. By the last of (A64), this requires $\Gamma_0 \to 0$, i.e. $Q \to 0$. In this limit we get

$$\Gamma_0 = \Delta_Q - \Delta_0 \simeq \Gamma_1 Q\,, \tag{A67}$$

which holds in the perceptron case. The saddle point equations therefore become equivalent to the one found in the perceptron.

### 6. Distribution of synaptic weights

The distribution of synaptic weights is:

$$P(W) = \left\langle\left\langle \frac{1}{\Omega} \int_0^\infty \prod_{li} dW_{li} \prod_{\mu=1}^{\alpha N} \Theta\left[ \frac{\sigma^\mu}{\sqrt{K}} \left( \sum_{l=1}^{K} c_l\, g\left( \sqrt{\frac{K}{N}} \sum_{i=1}^{N/K} W_{li} \xi_{li}^\mu - \sqrt{\frac{N}{K}} \theta d \right) - K\theta s \right) - \kappa \right] \delta(W - W_{11}) \right\rangle\right\rangle_{\{\xi^\mu, \sigma^\mu\}} \tag{A68}$$

with respect to the first weight of the first sub-perceptron $W_{11}$ for simplicity. Resorting to the replica method by introducing $n$ replicas we have:

$$P(W) = \lim_{n\to 0} \mathop{\mathbb{E}}_{\sigma^\mu} \int_0^\infty \prod_{i,l,a} dW_{li}^a \int \prod_{\mu,a,l} \frac{d\lambda_{l\mu}^a \, d\hat{\lambda}_{l\mu}^a}{2\pi} \, e^{i\lambda_{l\mu}^a \hat{\lambda}_{l\mu}^a} \prod_{\mu=1}^{\alpha N} \Theta\left[\frac{\sigma^\mu}{\sqrt{K}}\left(\sum_l c_l \, g(\lambda_{l\mu}^a) - K\theta s\right) - \kappa\right] \delta(W - W_{11}) \\ \times e^{i\sqrt{\frac{N}{K}}\theta d \sum_{\mu,a,l} \hat{\lambda}_{l\mu}^a} \prod_{l,i,\mu} \left\langle e^{-i\xi_{li}^\mu \sqrt{\frac{K}{N}} \sum_a W_{li}^a \hat{\lambda}_{l\mu}^a} \right\rangle_{\xi_{li}^\mu} \tag{A69}$$

Repeating the same steps as in section A 3 we have

$$P(W) = \lim_{n\to 0} \int \prod_{a<b,l} \frac{dq_l^{ab} \, d\hat{q}_l^{ab}}{2\pi K/N} \int \prod_{a,l} \frac{dQ_l^a \, d\hat{Q}_l^a}{2\pi K/N} \int \prod_{a,l} \frac{dM_l^a \, d\hat{M}_l^a}{2\pi\sqrt{K/N}} \, e^{-\frac{N}{K}\sum_{a<b,l} q_l^{ab}\hat{q}_l^{ab} - \frac{N}{K}\sum_{a,l} Q_l^a \hat{Q}_l^a - \frac{N}{K}\overline{W}\sum_{a,l}\hat{M}_l^a} \\ \times e^{N\alpha \, G_E(q_l^{ab}, Q_l^a, M_l^a) + (\frac{N}{K}-1)\, G_S(\hat{q}_l^{ab}, \hat{Q}_l^a, \hat{M}_l^a)} \int_0^\infty \prod_{l,a} dW_l^a \, \delta(W - W_1) \, e^{\sum_{a<b,l} \hat{q}_l^{ab} W_l^a W_l^b + \sum_{a,l} \hat{Q}_l^a (W_l^a)^2 + \sum_{a,l} \hat{M}_l^a W_l^a} \tag{A70}$$

where the entropic and energetic terms are the same as in Eqs. (A14a), (A14b) and the order parameters are those defined in (A11). In the limit $n \to 0$ therefore

$$P(W) = \lim_{n\to 0} \int_0^\infty \prod_{l,a} dW_l^a \, \delta(W - W_1) \, e^{\sum_{a<b,l} \hat{q}_l^{ab} W_l^a W_l^b + \sum_{a,l} \hat{Q}_l^a (W_l^a)^2 + \sum_{a,l} \hat{M}_l^a W_l^a} \tag{A71}$$

provided the order parameters satisfy the same saddle point equations as before. Under the RS ansatz expression (A71) becomes:

$$P(W) = \Theta(W) \int Dz \, \frac{e^{-\frac{1}{2}(\hat{q} - 2\hat{Q})W^2 + (\sqrt{\hat{q}}z + \hat{M})W}}{\int_0^\infty dW \, e^{-\frac{1}{2}(\hat{q} - 2\hat{Q})W^2 + (\sqrt{\hat{q}}z + \hat{M})W}} \\ = \Theta(W)\sqrt{\hat{q} - 2\hat{Q}} \, e^{-\frac{1}{2}(\hat{q} - 2\hat{Q})W^2 + \hat{M}W} \int Dz \, e^{\sqrt{\hat{q}}Wz} \frac{G\left(-\frac{\sqrt{\hat{q}}z + \hat{M}}{\sqrt{\hat{q} - 2\hat{Q}}}\right)}{H\left(-\frac{\sqrt{\hat{q}}z + \hat{M}}{\sqrt{\hat{q} - 2\hat{Q}}}\right)} \tag{A72}$$

Notice that the dependence of $P(W)$ on $K$ and on the activation function is not explicit, but is concealed inside the order parameters that clearly depend on them through the saddle point they have to satisfy. Notice also that for $\alpha = 0$ the synaptic weight satisfies an exponential distribution

$$P(W) = \Theta(W)\hat{M}e^{-\hat{M}W} = \Theta(W)\frac{f_{\text{in}}}{\theta_d} e^{-\frac{f_{\text{in}}}{\theta_d}W} \tag{A73}$$

This is to be expected, since at $\alpha = 0$ the only constrain that is required apart for the fact that the synapses are non-negative, is that their average is $\overline{W} = \frac{\theta_d}{f_{\text{in}}}$.

#### a. *Distribution of synaptic weights in the maximal storage limit*

In the critical capacity limit $\alpha \to \alpha_c$ the expression of the distribution of synaptic weight greatly simplifies. Using the scalings in (A54) we find

$$P(W) = \Theta(W) \, e^{-\frac{A}{2dq}W^2 - \frac{B\sqrt{C}}{dq}W} \sqrt{\frac{A}{dq}} \int Dz \, e^{\frac{\sqrt{C}}{dq}Wz} \left[G\left(\sqrt{\frac{C}{A}} \frac{z - B}{\sqrt{\hat{q} - 2\hat{Q}}}\right)\Theta(z - B) - \sqrt{\frac{C}{Adq}}(z - B)\Theta(B - z)\right] \tag{A74}$$

Using the identity

$$\int Dz\, e^{az}\,(z+b)\,\Theta(-b-z) = (a+b)\,e^{\frac{a^2}{2}}\,H(a+b) - e^{-ab}\,G(b) \tag{A75}$$

we obtain

$$P(W) = H(-B)\,\delta(W) + \frac{1}{\sqrt{2\pi}W_\star} e^{-\frac{(W+BW_\star)^2}{2W_\star^2}}\,\Theta(W) \tag{A76}$$

where $W_\star \equiv \frac{\sqrt{C}}{A}$. As showed in [5] in the one layer neuron model the synaptic weight distribution changes from being exponential to being a Gaussian plus a spike consisting to a fraction $H(-B)$ of "silent" weights at the critical capacity. Indeed as constraints due to the training set are added, more and more synapses tend to assume low weight. This is the case also in our two layer neuron model. It is interesting to note that both the distribution of synaptic weight at finite $\alpha$ (A72) and at critical capacity are in form exactly the same as the one derived in [5] for the one layer neuron model; the dependence on the non-linearity induced by the dendrites is actually implicit in the order parameters.

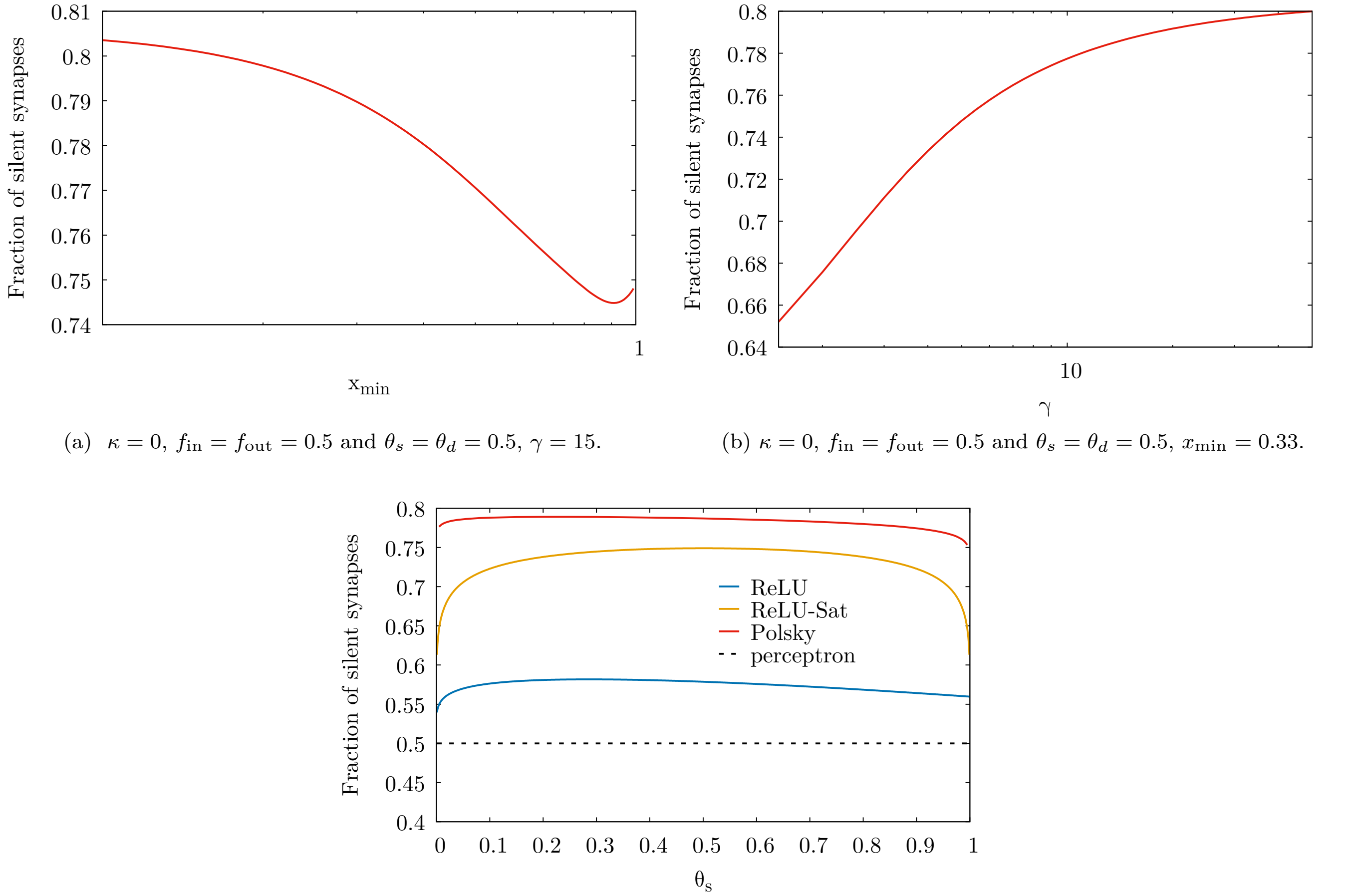


(a) $\kappa = 0$, $f_{\rm in} = f_{\rm out} = 0.5$ and $\theta_s = \theta_d = 0.5$, $\gamma = 15$.

(b) $\kappa = 0$, $f_{\rm in} = f_{\rm out} = 0.5$ and $\theta_s = \theta_d = 0.5$, $x_{\rm min} = 0.33$.

(c) $\kappa = 0$, $f_{\rm in} = f_{\rm out} = 0.5$ and $\theta_d = 0.5$, $x_{\rm min} = 0.33$, $\gamma = 15$.

Figure 2. In the top panels we plot the fraction of silent synapses for the Polsky non-linearity as a function the parameters $x_{\rm min}$ and $\gamma$. The bottom panel shows the fraction of silent synapses as a function of the somatic threshold, comparing the ReLU, the "saturating" ReLU and Polsky non-linearity when no robustness parameter $\kappa$ is imposed. The dashed black line represents the case of the one-layer neuron model, where the critical capacity $\alpha_c^{\rm perc} = 1$. In the captions of the panels we show the value of the fixed external parameters.

In Fig. **??** we show how the fraction of silent synapses $p_0 = H(-B)$ depends on the somatic threshold for the ReLU, ReLU-Sat and Polsky non-linearities. We also show in the Polsky case, how $p_0$ depends on the parameters defining the shape of the function itself, $x_{\rm min}$ and $\gamma$.

### Appendix B: Numerical experiments

#### 1. Choice of the thresholds

In this section, we outline a possible selection of thresholds $\theta_d$ and $\theta_s$, that can be utilized in numerical experiments and in the comparison with the analytical results. For SGD and the non-linear neuron case, as described in equation (A2), the first threshold $\theta_d$ is chosen to ensure that the pre-activations are distributed within the active range of the non-linear transfer function at initialization. The second threshold, $\theta_s$, is selected to maintain consistency between the output coding level and the ground-truth coding level. In contrast, for the linear neuron (perceptron) case, only a single threshold $\theta$ is involved, which determines the mean of the weights. By applying a suitable rescaling, this threshold can be arbitrarily fixed, while the cross-entropy parameter $\gamma_{\text{ce}}$ and the learning rate $\zeta$ must be appropriately chosen.

To achieve the desired output coding level $f_{\text{out}}$ during the initial forward pass, two factors have to be considered:

1. The distribution of pre-activations incoming to the hidden units, in order to choose the first threshold $\theta_d$.
2. The probability of generating either 1 or 0 after applying the Heaviside function to the neuron output. This probability is determined by the value of $\theta_d$ and the specific functional form of the non-linearity applied to the hidden units. By selecting the second threshold $\theta_s$ based on these factors, we can achieve the desired output coding level.

Considering random weights $W$, the pre-activations incoming to the hidden units for a given pattern $\xi$ are computed as follows:

$$a = \sqrt{\frac{K}{N}} \sum_i W_i \xi_i \tag{B1}$$

In the limit where $N, K \to \infty$ and $\frac{N}{K} \to \infty$, the distribution of these pre-activations converges to a Gaussian distribution by the central limit theorem (CLT). The mean $\mu$ and variance $\sigma^2$ of this Gaussian distribution depend on the input coding level $f_{\text{in}}$, the dendritic threshold $\theta_d$, and the initial weight distribution, which we assume to be uniform for simplicity. In our analysis, we consider the first threshold $\theta_d$ to be of order 1 ($\theta_d \sim O(1)$), and the weights $W_i$ are randomly drawn from a uniform distribution in the range $[0, W_M]$, where $W_M$ is also of order 1 ($W_M \sim O(1)$).

The mean and variance of the i.i.d. variables $\xi_i$ and $W_i$ are given by:

$$\mathbb{E}[\xi_i] = f_{\text{in}} \tag{B2}$$

$$\text{Var}[\xi_i] = f_{\text{in}}(1 - f_{\text{in}}) \tag{B3}$$

$$\mathbb{E}[W_i] = \frac{W_M}{2} \tag{B4}$$

$$\text{Var}[W_i] = \frac{W_M^2}{12} \tag{B5}$$

Since $\xi_i$ and $W_i$ are both i.i.d., the mean and variance of their product can be computed using the following properties:

$$\mathbb{E}[\xi_i W_i] = \mathbb{E}[\xi_i]\mathbb{E}[W_i] = f_{\text{in}} \frac{W_M}{2} \tag{B6}$$

$$\text{Var}[\xi_i W_i] = (\text{Var}[\xi_i] + \mathbb{E}^2[\xi_i])(\text{Var}[W_i] + \mathbb{E}^2[W_i]) - \mathbb{E}^2[\xi_i]\mathbb{E}^2[W_i] \tag{B7}$$

$$= \text{Var}[\xi_i](\text{Var}[W_i] + \mathbb{E}^2[W_i]) + \text{Var}[W_i]\mathbb{E}^2[\xi_i] \tag{B8}$$

$$= f_{\text{in}}(1 - f_{\text{in}}) \left( \frac{W_M^2}{12} + \frac{W_M^2}{4} \right) + \frac{W_M^2}{12} f_{\text{in}}^2 \tag{B9}$$

$$= \frac{1}{12}(4 - 3f_{\text{in}}) f_{\text{in}} W_M^2 \tag{B10}$$

To estimate the mean and standard deviation of the pre-activations incoming to the hidden units (B1), we can apply the central limit theorem (CLT) to the sum of the products $\xi_i W_i$:

$$\mathbb{E}[a] = \mathbb{E}\left[ \sqrt{\frac{K}{N}} \sum_{i=1}^{N/K} \xi_i W_i \right] = \sqrt{\frac{K}{N}} \sum_{i=1}^{N/K} \mathbb{E}[\xi_i W_i] = \sqrt{\frac{N}{K}} f_{\text{in}} \frac{W_M}{2} \tag{B11}$$

$$\text{Var}[a] = \text{Var}\left[\sqrt{\frac{K}{N}}\sum_{i=1}^{N/K}\xi_i W_i\right] = \frac{K}{N}\sum_{i=1}^{N/K}\text{Var}[\xi_i W_i] = \frac{1}{12}(4-3f_{\text{in}})f_{\text{in}}W_M^2 \tag{B12}$$

The mean of the Gaussian distribution of the preactivations is:

$$\mu = \mathbb{E}[a] - \theta_d\sqrt{\frac{N}{K}} = \sqrt{\frac{N}{K}}f_{\text{in}}\frac{w_M}{2} - \theta d\sqrt{\frac{N}{K}} \tag{B13}$$

It is reasonable (given the form of the transfer functions we consider) to center the Gaussian distribution of the pre-activations around 0, so that $\mu = 0$ and from (B13) we obtain:

$$W_M = \frac{2\theta_d}{f_{\text{in}}} \tag{B14}$$

that self-consistently gives $W_M \sim O(1)$ if $\theta_d \sim O(1)$.

The standard deviation of the Gaussian distribution of the pre-activations is:

$$\sigma = \sqrt{\text{Var}[a]} = \frac{W_M}{2}\sqrt{\frac{1}{3}(4-3f_{\text{in}})f_{\text{in}}} \tag{B15}$$

We can treat the standard deviation $\sigma$ as a parameter and solve for the first threshold $\theta_d$:

$$\theta_d = \sigma\sqrt{\frac{3f_{\text{in}}}{(4-3f_{\text{in}})}}. \tag{B16}$$

For example with $f_{\text{in}} = 0.5$ and by choosing $\sigma = 1$ for the Polsky transfer function we find $\theta_d = 0.775$. After determining the value of $\theta_d$, we can proceed to fix $\theta_s$. Let $h_k = g(a_k)$ denote the activations of the hidden units, where $g$ is the transfer function and $a_k$ is the pre-activation signal arriving at the dendritic unit $k$. Our goal is to estimate the probability of generating a 1 in the neuron's final output and ensure that it is equal to $f_{\text{out}}$, thereby preserving the output coding level on average. Given the application of a Heaviside transfer function to the output, this probability can be expressed as:

$$p\,(\text{out}=1) = p\left(\frac{1}{K}\sum_{k=1}^{K}h_k > \theta_s\right) \tag{B17}$$

$$= 1 - F_{\mathcal{N}(\mu_2,\sigma_2^2)}(\theta_s) = f_{\text{out}} \tag{B18}$$

where $F$ is the cumulative density function, and in (B18), we applied the CLT once more, which ensures that the output preactivations follow a Gaussian distribution with mean $\mu_2$ and standard deviation $\sigma_2$, implicitly dependent on the transfer function applied to the hidden units and their preactivation distribution.

It is therefore possible to find $\theta_s$ by solving the equation:

$$F_{\mathcal{N}(\mu_2,\sigma_2^2)}(\theta_s) = 1 - f_{\text{out}}. \tag{B19}$$

To define the values of $\mu_2$ and $\sigma_2$, we denote the hidden unit activations by $h_k = g(a_k)$ and recall that we imposed $P(a_k) = \mathcal{N}_{0,1}(a_k)$ when calculating the first threshold $\theta_d$. The CLT can also be applied to the activations $h_k$, yielding $P\left(\frac{1}{K}\sum_{k=1}^{K}h_k\right) = \mathcal{N}_{\mu_h,\sigma_h^2}$, where $\mu_h$ and $\sigma_h^2$ are the mean and variance of $h_k$, respectively. These values can be calculated for the Polsky transfer function as follows

$$\mathbb{E}[h_k] = \int_{-\infty}^{\infty} g(x)\mathcal{N}_{(0,1)}(x)dx = 0.369 \tag{B20}$$

$$\text{Var}[h_k] = \int_{-\infty}^{\infty} g^2(x)\mathcal{N}_{(0,1)}(x)dx - E^2[h_k] = 0.202 \tag{B21}$$

We see that for $f_{\text{out}} = 0.5$ we can simply impose:

$$\mathbb{E}[h_k] = \theta_s \tag{B22}$$

For Polsky transfer functions applied to the hidden units, we can numerically evaluate (B20), yielding $\theta_s = \mathbb{E}[h_k] = 0.369$.

Another notable case is that of Heaviside transfer functions, where $\mathbb{E}[h_k] = 0.5$. In this scenario, we can employ a simpler argument to estimate the thresholds: since the output of the hidden units is independent of $\theta_d$, the only free parameter is $\theta_s$. We have[2] $p\left(\sum_{k=1}^{K} h_k > K\theta_s\right) = 1 - \theta_s$. By imposing $1 - \theta_s = f_{\text{out}}$, we obtain $\theta_s = 1 - f_{\text{out}}$, which, for $f_{\text{out}} = 0.5$, is equivalent to (B22).

#### 2. Choice and scaling of the hyper-parameters

Consider the transfer function implemented by the neuron with non-linear dendritic branches, which represents the output preactivation prior to the thresholding operation performed by the $\Theta$-function:

$$\Delta^{\mu}_{\text{out}} = \frac{1}{\sqrt{K}} \sum_{l=1}^{K} c_l\, g\left(\sqrt{\frac{K}{N}} \sum_{i=1}^{N/K} W_{li}\xi^{\mu}_{li} - \sqrt{\frac{N}{K}}\theta_d\right) - \sqrt{K}\theta_s \tag{B23}$$

we impose that $W \in \left[0, \frac{2\theta_d}{f_{in}}\right]$ and consequently:

$$W \sim O\left(\frac{\theta_d}{f_{in}}\right).$$

If we assume that: $\theta_d \sim O(1)$, and $\theta_s \sim O(1)$ we have: $W \sim O\left(\frac{1}{f_{in}}\right)$.

We also have: $\sum_{i=1}^{N/K} W_{li}\xi^{\mu}_{li} \sim O\left(\frac{N}{K}\right)$. Consequently, from (B23), we observe that the dendritic pre-activations scale as:

$$\left(\sqrt{\frac{K}{N}} \sum_{i=1}^{N/K} W_{li}\xi^{\mu}_{li} - \sqrt{\frac{N}{K}}\theta_d\right) \sim O\left(\sqrt{\frac{N}{K}}\right). \tag{B24}$$

Consequently, the pre-activation variance remains finite in the limit $N \to \infty$, which is the primary motivation for choosing these scalings, as the pre-activation variance is a crucial factor in ensuring the consistency of the learning setting when varying the input dimensionality $N$ and the number of dendritic branches $K$.

Applying the same type of consideration to the pre-activation of the single output node in (B23) and recalling that $c_l = 1\ \forall\, l$ and $g(\cdot) \sim O(1)$, we find that it scales as:

$$\Delta^{\mu}_{\text{out}} \sim O\left(\theta_s\sqrt{K}\right) \tag{B25}$$

*Least-action learning algorithm (LAL)* In the LAL algorithm, when a wrong prediction occurs for a specific pattern, we identify the neurons that contribute to the error, i.e., the neurons at positions $l_*$ that have negative values for the following local stability measure over the dendritic branches:

$$\delta^{\mu}_{l} = -\sigma^{\mu}\left(\sqrt{\frac{K}{N}} \sum_{i=1}^{N/K} W_{li}\xi^{\mu}_{li} - \sqrt{\frac{N}{K}}\theta_d\right) \tag{B26}$$

and then we proceed to updating the weights with the following (possibly stochastic) rule:

$$w_{il_*} \leftarrow w_{il_*} + \zeta\sigma^{\mu}\xi^{\mu}_{il_*} \tag{B27}$$

By requiring an update of the same order as the weights, i.e., $w \sim O(\theta_d) \implies \zeta\sigma^{\mu}\xi^{\mu}_{il_*} \sim O(\theta_d)$, we obtain the desired scaling for the learning rate $\zeta$:

$$\zeta \sim O\left(\frac{\theta_d}{f_{in}}\right) \tag{B28}$$

---

[2] The probability of generating a 1 in the output is $\frac{\theta_s}{K}$. To preserve the output coding level, we set $\theta_s = K\,(1 - f_{\text{out}})$.

*Stochastic Gradient Descent (SGD) with cross-entropy loss* Recalling the expression for the cross-entropy loss used to investigate the performance of stochastic gradient descent (SGD) on the neuron:

$$\mathscr{L}_{ce}\left(\Delta^{\mu}_{\text{out}}\right) = \frac{1}{2\gamma_{ce}} \log\left(1 + \exp\left(-2\gamma_{ce}\Delta^{\mu}_{\text{out}}\right)\right) \tag{B29}$$

we can estimate the gradients of the cross-entropy loss, which are used in the SGD update, as follows:

$$\frac{\partial \mathscr{L}_{ce}\left(\Delta\right)}{\partial \Delta} = -\tfrac{1}{1+\exp(2\gamma_{ce}\Delta)} \tag{B30}$$

$$\sim O(1) \tag{B31}$$

where the last step holds (due to (B25)) if:

$$\gamma_{ce} \sim O\left(\frac{1}{\theta_s\sqrt{K}}\right). \tag{B32}$$

Proceeding with the derivative w.r.t. to the weights, we obtain:

$$\frac{\partial \mathscr{L}_{ce}\left(\Delta(W)\right)}{\partial W} = \frac{\partial \Delta(W)}{\partial W}\frac{\partial \mathscr{L}_{ce}\left(\Delta\right)}{\partial \Delta} \tag{B33}$$

$$= \frac{\partial}{\partial W}\left(\frac{1}{\sqrt{K}}\sum_{l=1}^{K} c_l\, g\left(\sqrt{\frac{K}{N}}\sum_{i=1}^{N/K} W_{li}\xi^{\mu}_{li} - \sqrt{\frac{N}{K}}\theta_d\right) - \sqrt{K}\theta_s\right)\frac{\partial \mathscr{L}_{ce}\left(\Delta\right)}{\partial \Delta} \tag{B34}$$

$$= \left[\frac{1}{\sqrt{K}}\sum_{l=1}^{K}\left(c_l\, g'\left(\sqrt{\frac{K}{N}}\sum_{i=1}^{N/K} W_{li}\xi^{\mu}_{li} - \sqrt{\frac{N}{K}}\theta_d\right)\sqrt{\frac{K}{N}}\sum_{i=1}^{N/K}\xi^{\mu}_{il}\right)\right]\frac{\partial \mathscr{L}_{ce}\left(\Delta\right)}{\partial \Delta} \tag{B35}$$

$$\sim O\left(\frac{1}{\sqrt{K}}\sqrt{\frac{K}{N}} f_{in} N\right) O(1) \tag{B36}$$

$$\sim O\left(f_{in}\sqrt{N}\right) \tag{B37}$$

where in the penultimate step, we have used the fact that the derivative of the function $g$ is bounded within the interval $[0,1]$ so that $g'(\cdot) \sim O(1)$; that $\sum_{i=1}^{N/K} \xi^{\mu}_{il} \sim O(f_{in}\frac{N}{K})$; and (B31).

At this point, recalling that the SGD update rule is:

$$w_{il} \leftarrow w_{il} - \zeta \nabla_{w_{il}} \mathscr{L}(w_{il}) \tag{B38}$$

and imposing an update of the same order as the weights, i.e., $w \sim O(\theta_d) \implies \zeta\nabla_w \mathscr{L}(w) \sim O(\theta_d)$, we find the desired scaling for the learning rate $\zeta$:

$$\zeta \sim O\left(\frac{\theta_d}{f_{in}\sqrt{N}}\right) \tag{B39}$$

*Comparison between the dendritic and the linear neuron* The transfer function of the linear neuron (i.e. the perceptron model), is given by:

$$\Delta^{\mu}_{\text{perc, out}} = \frac{1}{\sqrt{N}}\sum_{i=1}^{N} W_i \xi^{\mu}_i - \sqrt{N}\theta_s. \tag{B40}$$

From this expression, and observing that the weights scale with $\theta_s$, we obtain the following scalings for the learning rate $\zeta$ and cross-entropy parameter $\gamma_{ce}$:

$$\zeta_{LAL} \sim O\left(\frac{\theta_s}{f_{in}}\right), \qquad \zeta_{SGD} \sim O\left(\frac{\theta_s}{f_{in}\sqrt{N}}\right), \qquad \gamma_{ce} \sim O\left(\frac{1}{\theta_s\sqrt{N}}\right). \tag{B41}$$

---

[1] E. Gardner, Journal of Physics A: Mathematical and General **21**, 257 (1988).

[2] E. Gardner and B. Derrida, Journal of Physics A: Mathematical and General **21**, 271 (1988).
[3] M. Mezard, G. Parisi, and M. Virasoro, *Spin glass theory and beyond: An Introduction to the Replica Method and Its Applications*, Vol. 9 (World Scientific Publishing Company, 1987).
[4] A. Engel and C. Van den Broeck, *Statistical mechanics of learning* (Cambridge University Press, 2001).
[5] N. Brunel, V. Hakim, P. Isope, J.-P. Nadal, and B. Barbour, Neuron **43**, 745 (2004).
[6] N. Brunel, Nature neuroscience **19** (2016), 10.1038/nn.4286.
